# Predictive visuo-motor communication through neural oscillations

**DOI:** 10.1101/2020.07.28.224949

**Authors:** Alessandro Benedetto, Paola Binda, Mauro Costagli, Michela Tosetti, Maria Concetta Morrone

## Abstract

The mechanisms coordinating action and perception over time are poorly understood. The sensory cortex needs to prepare for upcoming changes contingent on action, and this requires temporally precise communication that takes into account the variable delays between sensory and motor processing. Several theorists^1,2^ have proposed synchronization of the endogenous oscillatory activity observed in most regions of the brain^3^ as the basis for an efficient and flexible communication protocol between distal brain areas^2,4^, a concept known as “communication through coherence”. Synchronization of endogenous oscillations^5,6^ occurs after a salient sensory stimulus, such as a flash or a sound^7–11^, and after a voluntary action^12–18^, and this impacts directly on perception, causing performance to oscillate rhythmically over time. Here we introduce a novel fMRI paradigm to probe the neural sources of oscillations, based on the concept of perturbative signals, which overcomes the low temporal resolution of BOLD signals. The assumption is that a synchronized endogenous rhythm will modulate cortical excitability rhythmically, which should be reflected in the BOLD responses to brief stimuli presented at different phases of the oscillation cycle. We record rhythmic oscillations of V1 BOLD synchronized by a simple voluntary action, in phase with behaviourally measured oscillations in visual sensitivity in the theta range. The functional connectivity between V1 and M1 also oscillates at the same rhythm. By demonstrating oscillatory temporal coupling between primary motor and sensory cortices, our results strongly implicate communication through coherence to achieve precise coordination and to encode sensory-motor timing.

## Results

### Oscillation of behavioural sensitivity

Participants initiated each trial with a voluntary action (keypress), causing two grating patches of slightly different spatial frequency to appear above and below fixation, after a variable delay. Figure 1 shows spatial frequency discrimination performance averaged across all participants (Figure S1 shows results for individual participants). Discrimination accuracy was not constant, but oscillated rhythmically over time, and is well fit by a sinusoidal function of 5.4 Hz (R^2^ = 0.43, p_*corrected*_ = 0.005: inset Figure 1B). The first minimum in accuracy occurred 70 ms after action onset, consistent with “motor-induced suppression”^19^. However, the suppression repeats periodically, generating an oscillatory pattern in the theta range, consistent with previous findings of behavioural oscillations synchronized with the onset of action^12,14,17^.

**Figure 1.**
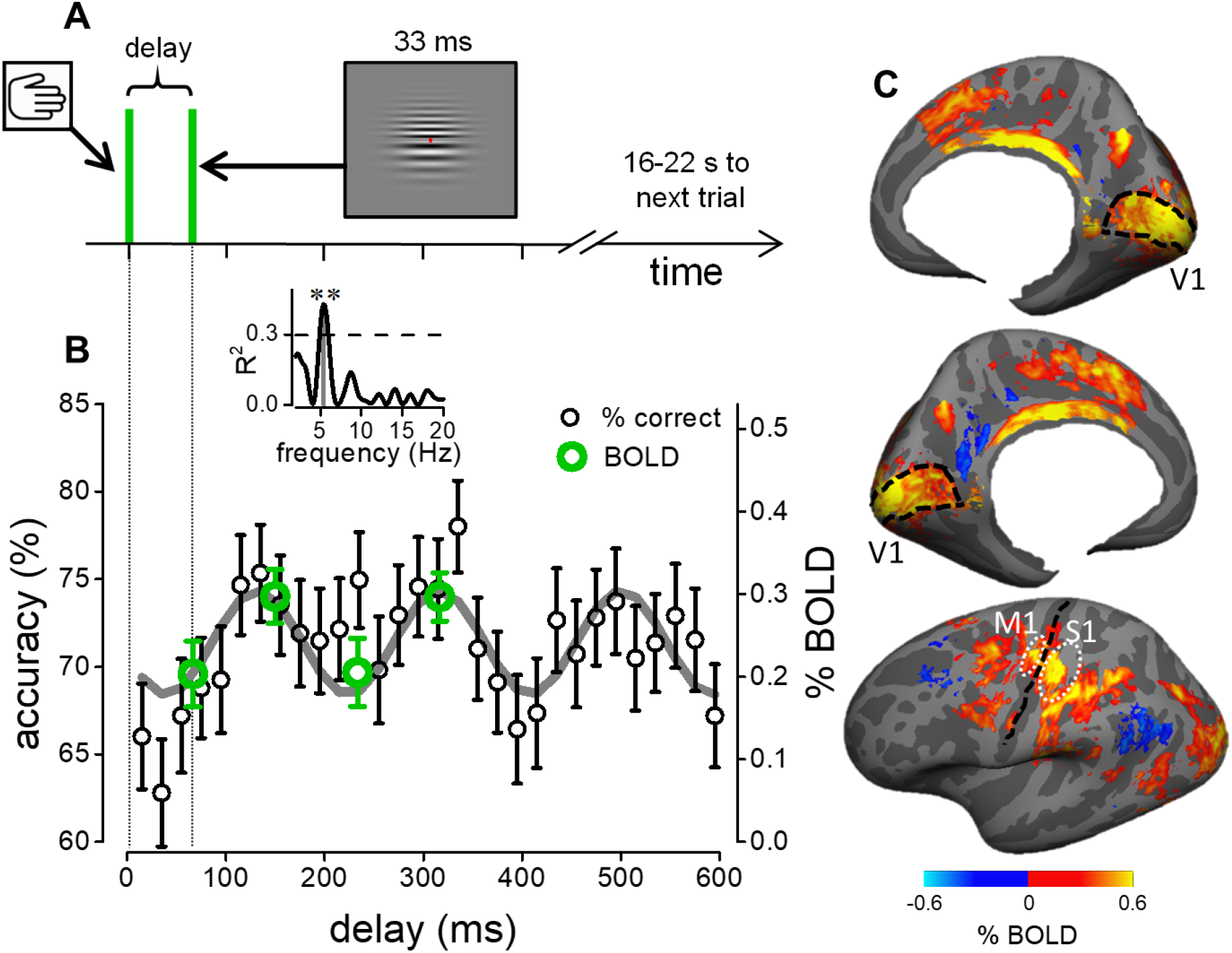
Experimental procedure, behavioural and BOLD responses. **A**: Schematics of the behavioural and the time-resolved fMRI design, measuring responses to visual stimuli following a voluntary action. The delay between stimulus presentation and action-onset varied randomly between 0 and 600 ms for the behavioural experiment, and randomly between four possible values (70, 150, 230 and 310 ms) for the fMRI experiment. **B**: Accuracy in the spatial frequency discrimination task as a function of visuo-motor delay (black symbols: mean ± s.e.m.; aggregate observer, N = 7), with the best sinusoidal fit of the accuracy time-course (grey line; 5.4 Hz). Green circles show % BOLD signal change in V1 (integral of the haemodynamic response from 3 to 12 s divided by time) at the four visuo-motor delays tested (extracted from Figure 2C: mean ± s.e.m.; N = 17). **Inset**: Goodness of the sinusoidal fit to the accuracy data in B as function of frequency, yielding a strong and significant peak at 5.4 Hz (R^2^ = 0.43, p_*correct*_ = 0.005,) above the 95th percentile of the R^2^ distribution obtained from fitting the permuted dataset with amplitude, phase and frequency as free parameter to obtain a corrected p value. **C**: Maps of the BOLD response to vision and action events (estimated at 6 s) projected on a template of the cortical surface and aggregated across N=17 participants; maps are masked at 0.05 significance after FDR correction. Black lines mark the central sulcus and V1 borders; white ovals mark the approximate surface location of the M1 and S1 ROIs.

### Oscillation of evoked BOLD responses in visual cortex

We repeated the psychophysical experiment of Figure 1A in an ultra-high magnetic field (7T) scanner, measuring the BOLD response evoked by a keypress followed by the visual grating. Given previous demonstrations that contrast sensitivity, known to reflect V1 activity, oscillates rhythmically after action onset^14^, we tested whether V1 activity is modulated by visuomotor delay by probing the peaks and the troughs of the accuracy oscillation. The BOLD response in V1 was strong and reliable, both when the stimuli were presented after the keypress action (*vision and action,* see activity map of Figure 1C), and when they were presented without the preceding keypress (*vision only*). However, the time-courses of the two responses were qualitatively different (Figure 2A) For stimuli not preceded by action, the BOLD response followed the typical V1 haemodynamic (Figure 2A, black symbols); for stimuli preceded by the keypress action, the BOLD response reached a similar peak, but attenuated more rapidly (Figure 2A, blue-green symbols). This difference cannot be accounted for by a single multiplicative or additive factor (see Model section below).

**Figure 2.**
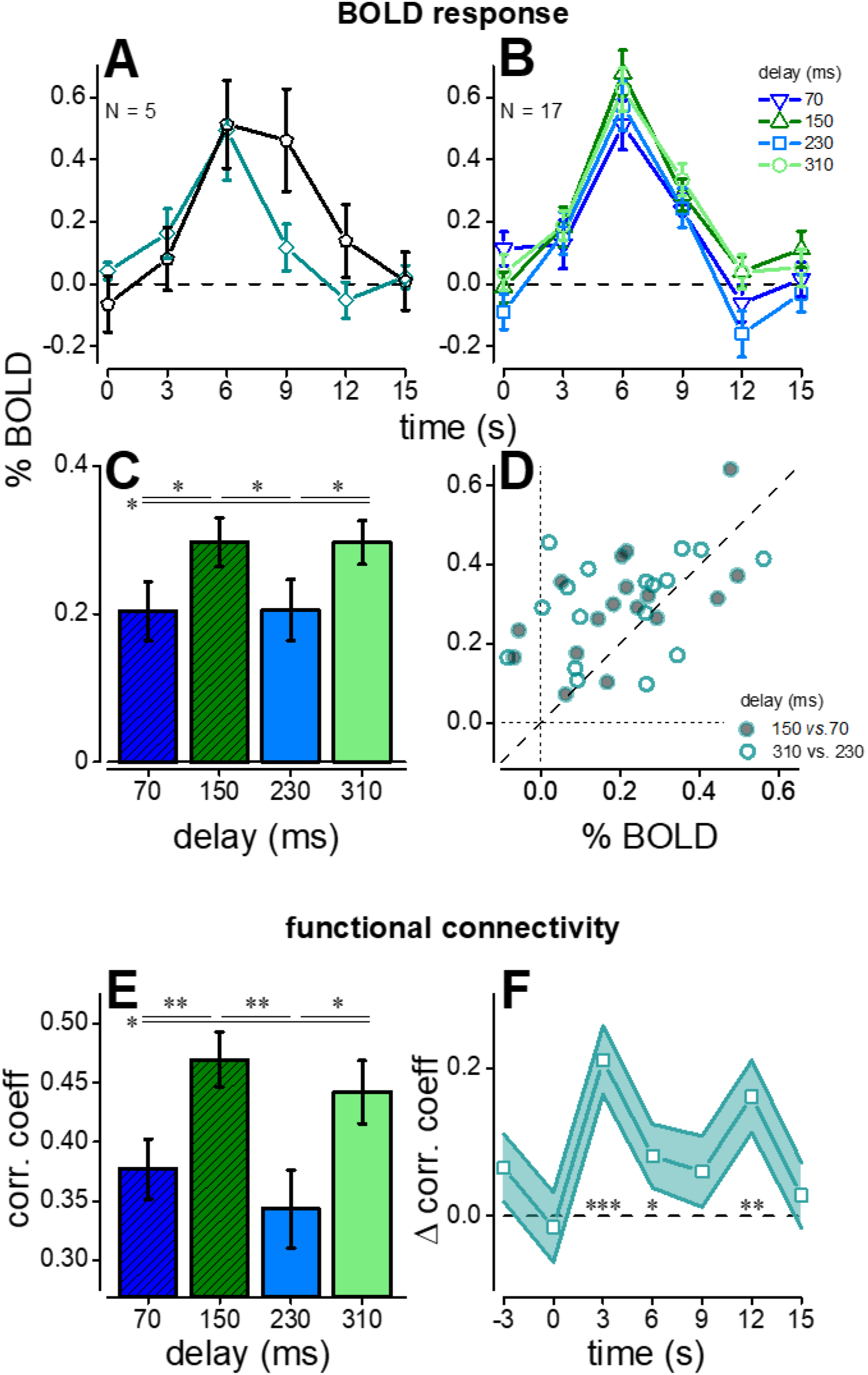
Oscillation of V1 BOLD responses and V1-M1 functional connectivity. **A**: BOLD response (estimated as GLM beta weights using the finite impulse-response function deconvolution approach) for the V1 subregion representing the stimulus area; means with s.e.m. from a subset of N=5 participants in the *vision only* (black curve) and *vision and action* condition (blue-green curve). **B**: Timecourse of the GLM beta weights (mean and s.e.m. for all N=17 participants) representing the BOLD response to *vision and action* events in the stimulated V1, separately for the 70, 150, 230 and 310 ms visuo-motor delays. **C**: Integral of the V1 response amplitude in the 3-15s interval, divided by time, as function of visuo-motor delay. Asterisks mark statistical significance (0.05 > * > 0.01) of post-hoc paired one-tailed t-tests (Bonferroni-Holm corrected for multiple comparisons) comparing pairs of visuo-motor delays: 70 ms vs 150 ms: t(16) = 2.88, p = 0.01; 150 ms vs 230 ms: t(16) = 2.54, p = 0.021; 230 ms vs 310 ms: t(16) = 2.20, p = 0.042; 70 ms vs 310 ms: t(16) = 2.33, p = 0.033). **D:** V1 response amplitude in individual participants, plotting delay 150 ms against 70 ms (filled symbols) or delay 310 ms against 230 ms (empty symbols). The large majority of points lie above the bisection of the axes (dashed line) implying that V1 BOLD responses were higher for *vision and action* events associated with the peak of psychophysical performance, compared to those associated with the minima of performance. **E**: M1-V1 functional connectivity, estimated as the correlation coefficient between residuals of the GLM fit (integral between −3 to 12 s divided by time) and plotted as a function of visuo-motor delay. Bars show mean and s.e.m of the bootstrapped aggregate observer. Asterisks (here and other panels) indicate statistical significance: ns > 0.05 > * > 0.01 > ** > 0.001 > ***. **F:** Time-course of the difference of the correlation coefficients of M1 and V1 residuals at the peak vs. trough delays at each TR around the keypress action. Symbols and error bars are mean and from the bootstrapped aggregate observer.

Figure 2B shows average haemodynamic responses separately for the four stimulus delays (relative to keypress). The responses to stimuli presented at delays yielding minimal psychophysical accuracy (blue curves) were clearly less than to those at delays yielding maximal psychophysical accuracy (green curves). Figure 2C shows the average of the response over the interval of 3 to 12 s (integral divided by time). Responses at delays of 150 and 310 ms are significantly greater than at 70 and 230 ms. Repeated measures ANOVA showed a significant effect of stimulus delay (F(3,48) = 4.17; p = 0.01; η^2^ = 0.21), with amplitude at 70 and 230 ms significantly lower than the other two. The effect occurred in most of the 17 participants (Figure 2D). The BOLD rhythmic modulation between the combined peak versus the combined trough delays remains significant 6 s after action onset (Figure S2 C, continuous line). In this analysis the BOLD regressors were aligned to the action onset, which produces massive responses in many motor and visual brain areas. However, similar visual BOLD modulations occurred when aligning the response to stimulus onset (Figure S2, A, B and Figure S2C, dashed line).

The V1 BOLD response amplitudes (Figure 1B, green circles) are highly consistent with the behavioural data collected outside the scanner, with the minima and the maxima of BOLD responses coinciding with the minima and maxima of performance in the spatial frequency discrimination task, confirming our hypothesis of a rhythmic modulation of V1 excitability.

### BOLD correlations in the sensory-motor network

Having established that BOLD responses in V1 modulate rhythmically at around 5 Hz, we studied whether a similar oscillation also occurs in motor or somatosensory areas. For each participant, we defined ROIs in motor (M1) and somatosensory (S1) primary cortex using independent fMRI acquisitions and measuring the response to keypress used to report the perceptual decision (see ROIs in Supplementary Figure S1 D). The haemodynamic responses of M1 and S1 to *vision and action* events are different from those in V1 (compare panel A to panels D&G in Figure S2): the response occurred earlier (reflecting activity during action preparation^20^), and had a more pronounced negative lobe. However, the response was not modulated by the visuo-motor delay (Figure S2, panels E&H), at any time of the haemodynamic response (Figure S2, panels F&I), contrary to reported evidence when reaction times are also measured^21^.

The absence of M1 BOLD modulation with visual delay is consistent with the fact that motor cortex does not activate in response to visual stimuli. However, synchronous oscillatory activity may be present in M1 and shared with V1^15^. We used a correlation analysis to investigate this possibility. We subtracted the responses estimated independently at each time-point, and measured the shared trial-to-trial variability of BOLD residual signals between areas V1 and M1, obtaining a measure of functional connectivity. Figure 2E shows the correlation coefficients for the aggregate subject at each visuomotor delay; the correlation has an oscillatory pattern, with stronger correlations at delays of 150 and 310 ms. Similarly, a pairwise t-test comparing correlations at the peak and trough delays across participants is statistically significant (t(16) = 1.80, p = 0.044). The correlation is strong and significant at all times (bootstrap of all trials of the aggregate subject: p < 0.001; one-sample t-test across participants: all t(16) > 3.5, p < 0.003); however, and importantly, there was no modulation with delay for times preceding stimuli onset. The strongest increase in correlation occurred at +3 and +12 s, being about 20% (Figure 2F). This indicates that the increased correlations are not due to incomplete deconvolution of the signal, given that it is absent at 0 s and does not follow the haemodynamic response. Interestingly, it is strong at 12 s after onset, where the BOLD responses differ also in sign for the various delays. Coupling between M1 to V1 is enhanced precisely at delays corresponding to maximum stimulus discriminability and BOLD V1 cortical excitability.

### Retinotopic dependence of rhythmic oscillation

The contrast map between BOLD responses for peak and trough delays (Figure S1 E) shows that many areas of the occipital pole are modulates by visuo-motor delay. Consistent with the transient nature of the stimuli, BOLD activity at all delays is particularly strong in the V1/V2 region, with additional foci of activity in V3, V4 and parieto-occipital sulci (Figure 3A). It is clear on inspection that the effect of delay is prominent in V1, with BOLD responses at 150 ms and 310 ms delays stronger and more diffuse than at 70 ms or 230 ms. The effect of delay occurs at all V1 eccentricities (Figure 3B), even beyond the area covered by the stimulus (the bottom icon shows stimulus contrast across eccentricity). This implies that the effect of delay is independent of the stimulus characteristics, and probably driven by automatic and pervasive modulation of the excitability of the entire early visual cortex, which oscillates rhythmically in synchrony with action onset (see Model Section).

**Figure 3.**
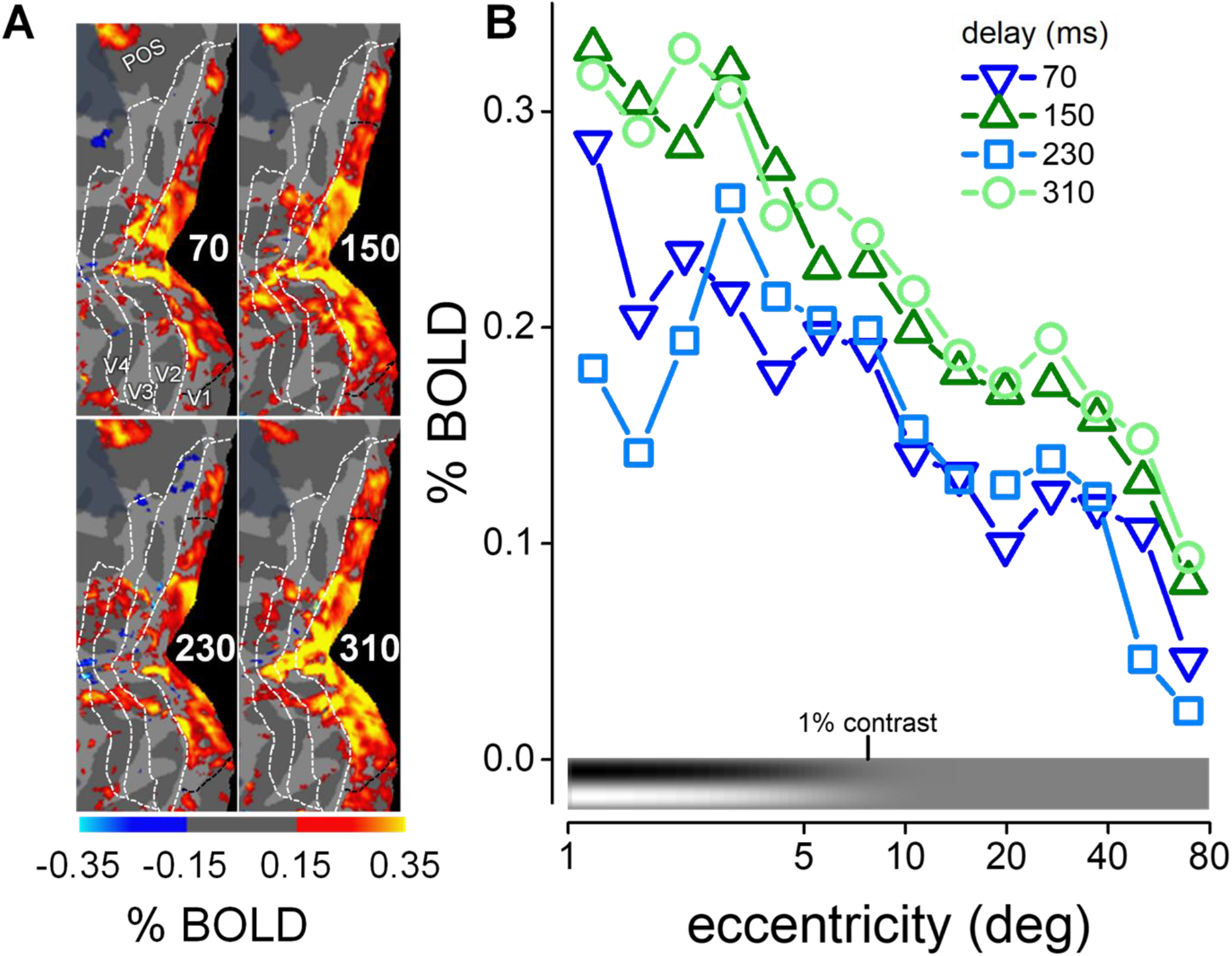
Effect of eccentricity on BOLD oscillations. **A**: Maps of the posterior cortex response to *vision and action* events with the four visuomotor delays. BOLD responses were computed from the aggregate observer after projecting functional data from both hemispheres onto the mirror-symmetric template of the cortical surface. Maps are masked by the value of beta-weights. White dashed lines indicate the borders between V1, V2, V3 and V4; dark dashed lines delimit the part of V1 stimulated by the gratings. The left panels show activity for the 70 and 230 ms visuo-motor delays, and the right panels for the 150 and 310 ms delays. **B**: V1 BOLD response (aggregate data, as shown in panel A maps) plotted as function of eccentricity in 14 non-overlapping logarithmic steps from 1 to 80 deg. The icon by the x-axis represents the variation of stimulus contrast over eccentricity.

### Model of the BOLD visuo-motor response

The overall pattern of results strongly suggests that visual and motor cortices become synchronized during a simple visuo-motor task (keypress followed by a visual stimulus), and that the BOLD amplitude oscillates rhythmically with visuo-motor delay, suggesting that both cortices are driven by synchronization of endogenous oscillations. The salient findings are:

1. BOLD responses in V1 have a different time course when associated with a voluntary action (Figure 2A).
2. The change of BOLD response with delay is not only multiplicative but includes an additive negative component (Figure 2B).

To capture both these effects, we developed a toy model simulating some well-known physiological properties of the visuo-motor loop, including corollary discharge signals during motor preparation that modulate V1, oscillatory rhythms and gain modulation of visual responses (see Figure 4). The two key hypotheses are: 1) V1 activity is modulated in synchrony with motor signals, both before and after action, through phase synchronization of endogenous rhythms at theta frequency; 2) the oscillation modulates V1 neuronal response gain.

**Figure 4.**
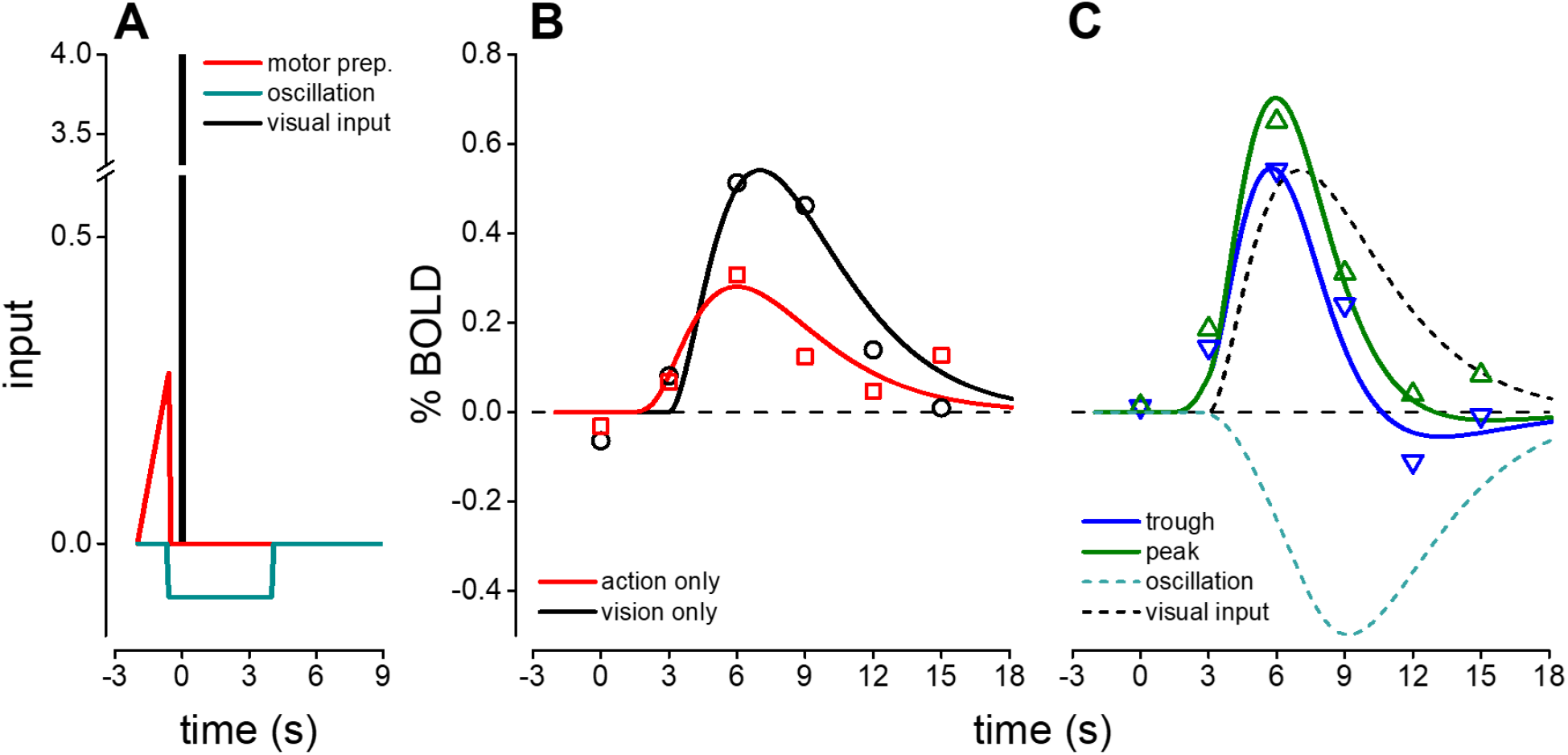
Simulation of V1 BOLD responses. **A:** The model assumed three components: motor preparation modelled with a ramp function (in red, starting −2 s before and completed by −550 ms before action-onset), neural oscillation as a negative boxcar (blue-green line, starting at −550 ms and completed 4 s after action onset), and visual input as a delta function (black line). Simulated curves in B-C were obtained by convolving the appropriately scaled inputs with a standard haemodynamic function^22^ with decay time τ = 2 s, order = 3 and delay δ = 3 s. **B-C:** simulated (lines) and observed (symbols) V1 BOLD responses to visual events presented without keypress (*vision only*, black curve in B and C), to keypress-only events (red curve in B and C), and to *vision and action* events associated with the trough (blue curve in C) or peak delays (green curve in C) of psychophysical performance.

We modelled the visual input with a delta function with amplitude chosen to maximize goodness of fit of V1 BOLD response to *vision-only* events, assuming a standard haemodynamic response^22^ (black trace in the inset, R^2^ = 0.91). To model the time-course of the V1 BOLD response to *vision and action* events, we assumed that the visual input is combined with a motor preparation signal given by the “readiness potential”, represented as a ramp signal starting 2 s before the action onset^23,24^. This readiness potential can also be recorded in V1 in the absence of visual stimulation (red curve in Figure 4B reporting the average activity in V1 in response to *action-only* events, see also maps in Figure S1 D), consistent with previous results^25–28^. Ramp amplitude in V1 was determined by the best fit of this response (R^2^ = 0.73, Amp=0.27). As schematically illustrated in Figure S3 A, the readiness potential is associated with (or may induce) network oscillations at theta frequencies. The synchronisation induces a negative and persistent signal, which we simulate over 4 s. The assumption of a negative signal is based on previous work showing that negative BOLD is associated with synchronized endogenous rhythms^29,30^, and with direct modelling of LFP and spiking activity associated with system dynamics^31,32^. The response to this negative signal, when summed with the responses to the motor preparation, predicted the M1 response (Figure S3 D), and when summed with the visual stimulus predicted the time course of the response to *vision and action* events for delays at the troughs of the oscillation (Figure 4C, blue curve; R^2^ = 0.91).

Finally, we postulated a small modulation of the gain of the visual response (estimated from the *vision only* condition) following the theta rhythm after action-onset (see Figure S3 A). This was achieved by moderate increase of the amplitude of the delta function that represents the visual input for the peak delay. A gain increase of about 33% is sufficient to fit the response to *vision and action* events for delays at the peak of the oscillation (Figure 4C, green trace; R^2^ = 0.92). Both the subtractive signal and the gain modulation are consistent with neuronal mechanisms associated to the phase of synchronized oscillations in sensory cortex^32^.

With very few parameters, the model captures quantitatively the many different aspects of V1 and M1 BOLD modulations. The functional connectivity result (Figure 2E–F) can also be explained by synchronization of activity and common signals shared by M1 and V1 (Figure S3 A), with their associated noise. The multiplicative response gain in V1 associated with the theta oscillations will contribute non-linearly to the co-variance between V1 and M1^33^. Numerical simulations (see caption of Figure S3) support this explanation.

## Discussion

Preparation of voluntary actions, such as finger-tapping, button-press, reaching, grasping and even isometric muscle contraction, can modulate the perception of visual stimuli^34^. From hundreds of milliseconds before action onset (during motor preparation) to up to 1 second after action offset, visual sensitivity oscillates within the theta range, phase-reset by action^12,14,17^. Here we found that V1 BOLD responses also oscillate at about 5 Hz following a voluntary action, strongly implicating V1 as the origin of the perceptual oscillation.

Although our paradigm did not allow us to correlate single trial performance with BOLD responses to directly assess the link between BOLD oscillation and performance, our data, together with previous results from our laboratory^11,12,14–17^, show that the aggregate oscillations are consistent across the population. Furthermore, it is known that ERP during motor preparation can predict individual oscillatory performance^15^. Interestingly, the resolution of the delay-dependent BOLD modulation was finer than that required for humans to judge simultaneity between action-onset and a visual stimulus (around 150 ms^35^), excluding cognitive effects. The BOLD V1 oscillation is also consistent with recordings in monkeys showing rhythmic theta-range modulation in V1^36–38^.

Our results provide strong evidence that voluntary action modulates V1 activity: not only was the visual response after a voluntary action different from that recorded during the *vision only* condition, but a voluntary action on its own, unaccompanied by any visual stimulus, also elicited a significant V1 BOLD response. The observed pattern of V1 responses is consistent with the models of active sensing or embodied perception, which propose that processing of sensory inputs is profoundly altered when input is actively sought through eye-, hand-, or body-movements, compared with passive stimulation^39,40^. Another well-described phenomenon associated with active sensing is motor-induced suppression of sensitivity, typically occurring in the first 100 ms after action onset. The phenomenon is very obvious after saccadic eye movements^41^, but also occurs after hand movements^42,43^, and can cause a measurable change in V1 BOLD response^26^. The attenuation of the response to stimulus presentation that we observe at 70 ms delay might be interpreted as the consequence of this motor-inhibition. However, if this modulation of the haemodynamic response were simply transient motor-suppression, we would have expected a monotonic increase with larger delays, rather than the rhythmic modulation we observe. Rather than a simple transient suppression, our results point to a cyclic alternation starting before action-onset. On this view, the function of the modulation is not to eliminate unstable sensory signals when sensors move, but rather is part of a cyclic mechanism of temporal binding^1,2,4,34^. The observed oscillation of BOLD responses and functional connectivity can be explained by synchronization of activity in motor and visual cortex by a common oscillatory rhythm present during the preparation phase. The anatomical pathways mediating the synchronization are at present unknown; they may not require a direct input of M1 to V1 and they may involve subcortical, e.g. thalamic, relays^44,45^.

We modelled all the complex features of our results, including the perceptual modulation, by assuming that synchronous oscillations cause a general subtractive effect on BOLD signal in V1^29–31^, and that cortical excitability undergoes a multiplicative oscillatory gain change, which follows the phase of the theta rhythm^32,36,37,46–48^. Both our key assumptions are consistent with the BOLD literature^49^. Our data and model framework exploit a mechanism of “communication through coherence”^2^ for transferring information between the sensory and the motor systems. This protocol is assumed to operate between distal cortical areas by synchronization of neural activity in local neuronal assemblies. Synchronized activity enhances effective connectivity within a local assembly and selectively improves the communication between assemblies^2,4^. Although synchronization typically occurs in the gamma-range (30-90 Hz)^1,4^, slower oscillations in the alpha and theta ranges can modulate the gamma rhythm and thereby affect communication^36,37,46–48,50^.

Other examples of communication through coherence exist in sensory-motor systems. Corticospinal gamma-band coherence occurs during movement preparation, and the coherence correlates with reaction times^51^. Functional connectivity between cortical areas changes, depending on the local phase of the endogenous theta oscillation in humans^52^. Theta range synchronous modulation has been interpreted as a multiplexing mechanism to allocate processing resources to sensory and motor functions^53^. As with most multiplexing communication systems, this oscillatory communication may be efficient and easy to time-lock. Our results corroborate these findings, and together suggest an efficient, predictive, and flexible communication system through synchronization of endogenous rhythms for sensory-motor coordination. Flexibility is a critical issue, given that sensory-motor synchronization needs to rapidly recalibrate to accommodate internal and external changes (for example, fatigue can affect motor delays, and illumination visual delays). This complex task may become more manageable if time is encoded in a cyclical function, represented within primary sensory areas in the form of periodic modulations of cortical excitability.

Several studies propose that the phase of theta oscillations drives selective and spatial attention^7,9,10,54–57^, which are well known to modulate low-level areas such as V1^58^. Attention may in principle contribute to our results, but this seems unlikely for several reasons. Firstly, the oscillation of V1 BOLD responses was not limited to the central visual field modulated by the stimulus, but extended to the far periphery of V1, which was neither stimulated nor task-relevant. Secondly, V1 representations of upper and lower halves of the stimulus oscillated in-phase, consistent with evidence that oscillations synchronized by voluntary actions have the same phase across multiple locations^12^, whereas attentional fluctuations typically have opposite phases across locations^7,9^. Thirdly, the V1 modulation was not stronger in the lower visual field, despite the lower-hemifield bias of attentional processes^59^. Another possibility is that oscillations of the rate of microsaccades or drift eye movements synchronized with keypress drive the rhythmic modulations reported here^60,61^. While we cannot dismiss this possible influence, in another study we measured microsaccade rate and found no correlation with the behavioural oscillation^16^.

Our results demonstrate that visuo-motor interactions have high temporal specificity, since perceptually indistinguishable changes in the delay between motor and visual events are sufficient to produce dramatic changes in the pattern of V1 activity, as well as of M1-V1 functional connectivity. Moreover, premotor signals (readiness potential) actively participate in the temporal recalibration of cortical activity necessary after adaptation to altered visuo-motor temporal delays for efficient processing^62^. All these examples may be interpreted as evidence for predictive coding of the future consequence of the action^1,42,43,63–68^ elaborated during motor preparation and represented within sensory cortex. These predictions may be instrumental in stabilizing perception across movements, as well as in endowering a sense of agency.

## Conclusion

The M1-V1 sensory-motor loop rhythmically oscillates at theta rhythm after a voluntary action, causing perceptual consequence such as modulation of visual sensitivity. The synchronized network dynamics may be instrumental for temporal binding in the visuo-motor control system. This mechanism could underly communication of the time of action from the motor cortex to the sensory cortex, and encode it with high precision over the extended period when relevant sensory signals may arise.

## Acknowledgments

This project was supported by the European Research Council (ERC) under the European Union’s Horizon 2020 program (Grant Agreement No 832813-GENPERCEPT and No 801715-PUPILTRAITS) and by MIUR - PRIN 2017 - grant 2017SBCPZY_02. We thank Eleanor Reynolds and David Burr for proofreading the manuscript. The authors would like to thank Dr. Brian Burns (GE Healthcare) for his implementation of the 3D-MP2RAGE sequence on the MR950 system.

## Author contributions

Conceptualization, A.B, P.B., and M.C.M.; Methodology, all authors. Investigation, A.B, M.C.M. and P.B.; Writing, all authors.

## STAR Methods

### RESOURCE AVAILABILITY

#### Lead Contact

Further information and requests for resources should be directed to the Lead Contact, Maria Concetta Morrone.

#### Materials Availability

This study did not generate new unique reagents.

#### Data and Code Availability

Analyses were performed with Freesurfer v6.0.0, SPM12, BrainVoyager 20.6, and FSL (version 5.0.10) packages And MATLAB (version R2020b). Matrices of the results are available at 10.5281/zenodo.4700266.

## EXPERIMENTAL MODEL AND SUBJECT DETAILS

### Human participants

Experimental procedures are in line with the declaration of Helsinki and were approved by the regional ethics committee (*Comitato Etico Pediatrico Regionale—Azienda Ospedaliero-Universitaria Meyer*—Firenze (FI)) and by the Italian Ministry of Health, under the protocol ‘*Plasticità e multimodalità delle prime aree visive: studio in risonanza magnetica a campo ultra-alto (7T)*’ version #1 dated 11/11/2015. Written informed consent was obtained from each participant, which included consent to process and preserve the data and publish them in anonymous form.

We recruited seven participants for the behavioural experiment (26-36 years old, mean age = 30, 3 females and 4 males), with normal or corrected to normal vision. Twenty-four healthy participants with normal or corrected-to-normal vision were recruited for the fMRI experiment. Three of them did not complete the scanning session and were excluded from the analyses. Of the remaining 21 participants (24-63 years old, mean age = 30, 9 females and 12 males), 17 took part in the main experiment that included 60±17 visual presentations over 5±2.5 fMRI acquisition series (mean ± standard deviation).

## METHOD DETAILS

### Behavioural experiment: setup and procedure

Behavioural measurements were made in a quiet room in dim light condition. The stimuli were generated in MATLAB (MATLAB r2010a; MathWorks), displayed with a graphics board ViSaGe (Cambridge Research System) on a linearized CRT monitor (Barco Calibrator 40×30 cm, resolution of 800×600 pxl, refresh rate 100 Hz, mean luminance 25 cd/m^2^) at 57 cm from the observer. Keypress responses were acquired by CB6 response box (Cambridge Research System) with a negligible estimated time error < 1ms (as per manufacturer’s specifications). Stimulus presentation was synchronized with the monitor framerate, implying that visuo-motor synchronization could be determined with an error between −10 and +0 ms.

The visual stimulus comprised two horizontal gratings (10% contrast, spatial frequency 1 c/º and 1.1 1 c/º) presented for 30 ms (3 frames) in the upper and lower visual field, both vignetted by a Gaussian window (σ = 5 deg) at screen centre (see inset Figure 1). The two gratings were separated by a Gaussian horizontal blank with space constant of 0.5 deg (σ). The two gratings were presented randomly between to top and the bottom field with each trial. The phase of each grating also varied randomly across trials. Participants maintained fixation on a small red dot (0.15 deg) at the centre of the display and pressed a key to start the trial (with the fourth finger of the right hand). The visual stimulus was displayed at 60 possible stimulus delays (corresponding from 1 to 60 monitor frames) from button press, chosen randomly on each trial from a uniform distribution between 0 and 600 ms in steps of 10 ms. The participant’s task was to indicate which grating (upper or lower) had the higher spatial frequency (i.e. a 2AFC spatial frequency discrimination task) by pressing a button with the index or the middle finger of the right hand.

Participants were instructed to wait at least 1 s between two successive keypresses (this interval was 1.6 s on average, with 1 s standard deviation). Trials with an ITI of less than 1 s (1.3% of trials) were excluded from further analysis. We collected on average 1008±187 trials for each participant.

### Behavioural oscillation analysis

To test whether visual accuracy fluctuated rhythmically over time, we pooled all trials from all participants (aggregate subject), sorted them in terms of visuo-motor delays and grouped them into 20 ms non-overlapping bins (but note the duration of the stimulus, 30ms). For each bin, we computed the proportion of correct responses and fitted the resulting time series with the following sine function:

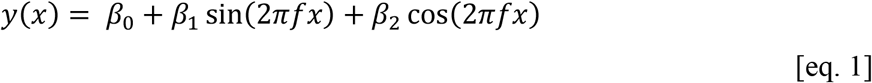

where ƒ represents the frequency of the oscillation and can vary between 2 and 20 Hz in steps of 0.1 Hz; *β*_0,_ *β*_1_ and *β*_2_ are free parameters to be estimated. Statistical significance was evaluated by comparing the goodness of fit (R^2^) for the original dataset, with the distribution of goodness of fits obtained on permuted datasets created by shuffling the original responses (10000 permutations). For each permutation, we selected the best fitting sinusoidal model, with frequency, amplitude and phase free parameters and compared the distribution of the obtained R2 with the R^2^ of the real data best fit. To use all free parameters for the fit of surrogate data allows to correct automatically for multiple comparisons of the frequency range.

To evaluate the consistency of amplitude and phase across subjects, we ran multivariable generalized linear model (GLM) applied to single-trial response^15,69^. For each participant, we fitted a linear regression model including as predictors a sine and a cosine for a given frequency of interest (between 3.5 and 10 Hz, resolution of 0.1 Hz). The fixed-effect linear regression parameters were estimated using standard least square method (LSM). The beta coefficients of the participants were tested against 0 by means of the bivariate Hotelling’s T-squared statistic, i.e. an extension of the Student’s t test to the multivariate domain.

### fMRI experiment: setup and procedures

Visual stimuli were presented by using a magnetic resonance-compatible goggle set (VisuaStimDigital, Resonance Technologies Inc., Northridge, California) with visual field of approximately 32×24 deg, 800×600 resolution, refresh rate 60 Hz (16.7 ms frame), mean luminance 25 cd/m^2^. Keypresses were recorded through a magnetic resonance-compatible response-box (Evoke Response Pad System, Resonance Technologies Inc., Northridge, California), positioned on the right hand of volunteers. The time of the keypress was recorded with a constant error of 10±0.5 ms, (due to the MS Windows software) and associated with the time-stamp of the frame in which it occurred.

The same visual stimulus and procedure of the behavioural experiment were used in the fMRI scans, with the following differences. To accommodate for the different refresh rate of the monitor, the duration of the stimulus was limited to two frames (33 ms). The high sensitivity of 7T fMRI allowed for recording reliable BOLD responses to such brief stimulus, even for small numbers of events. Stimulus presentations were synchronized with the monitor framerate, implying that visuo-motor synchronization could be determined with an error between −10 and +6 ms (slightly asymmetric due to the constant delay of the keypress time-stamp). As for the behavioural experiment, participants started the trial by pressing a key using the fourth finger (chosen based on the high sensitivity of the key). They were instructed to wait at least 15 s between two successive keypresses. The subject practice the task before entering the scanner till they reach a correct timing pace without using an internal count-down.

In the main experiment, the task was to silently perform the spatial-frequency discrimination task and to report an estimate of the average proportion of “up” responses at the end of each acquisition series.

In a small set of runs (not included in the main analyses), all 21 participants were required to report perceptual decisions with a second keypress, delayed by several seconds (15 s on average, with 4 s standard deviation) to minimize interference with the visual presentation. These trials were used to assess stimulus discriminability in the scanner and to check that performance was not saturated at 100% correct or at 50% correct. These trials were used to assess stimulus discriminability in the scanner; while most participants performed well above chance with an average 62±3% correct, few of them (5 out 17) complained that maintaining an accurate memory of the choice for so long time was very hard and performed near chance (below 54%). Excluding these participants brought average performance to 69%, very similar as in the behavioural experiment conducted outside the scanner (t(11) = 0.56, p = 0.581, considering the full population the t-test was t(16) = 2.45, p = 0.027). The paucity of trials across all participants did not allow for tracking oscillations in accuracy as function of visuo-motor asynchrony; this also prevented us from analysing the single-trial correlation between BOLD and behavioural responses.

For each participant we used the BOLD modulation associated with the decision keypress to locate cortical areas holding a representation of the keypress action both in M1 and S1 (see supplementary Figure S1). Primary visual cortex V1 was localized by separate retinotopic mapping data (two 45 deg wedges centred around the horizontal or vertical meridian, presented alternately for 5 TRs each, with no blanks, and six repetitions in total for a total of 120 TRs).

Data for the main experiment (with no keypresses for perceptual decisions) were collected in 17 of the 21 subjects. Visual stimuli were presented every 19 s on average (3 s standard deviation) at 4 asynchronies between the keypress and the visual stimulus (delay = 70, 150, 230, or 310 ms, with the −10:+6 error range indicated above); keypresses (and the following visual presentations) could not be aligned with the TR since participants pressed they key at their own pace; each acquisition series lasted 130 TR of 3s. The average number of trials (± 1 standard deviation) per participant were: 17±6 for delay 70 ms; 17±6 for delay 150 ms; 13±5 for delay 230 ms; 13±4 for delays 310 ms (privileging shorter delays that were perceptually indistinguishable^35^). In order to define the design matrix used for the GLM analyses, we assigned each *vision and action* event to the TR preceding action onset. This meant that in about 3% of trials, the visual presentation actually occurred 1TR later than indicated in the design matrix (this happened in about 5% of trials for the 310 ms delay, and 4%, 3%, and 2% for the 230, 150 and 70 ms delays respectively). We verified that aligning the design matrix to the visual stimulus rather than the action onset did not change the results (Figure S2, panel A-B). We also verified that the temporal distribution of events was homogeneous in the TR, using circular statistics (Rayleigh test for 70 ms delays: p = 0.34; 150 ms delays: p = 0.55; 230 ms: p = 0.89; 310 ms delays: p = 0.45). The results ensured that there were no temporal biases between conditions and therefore that event timing could not explain response amplitude differences. Given the limitations in scanning time at 7T (about 1h per subject per 6 months), testing was limited to 4 delays that matched the maxima and minima of psychophysical performance; this logically corresponds to testing the hypothesis that BOLD responses had the same oscillations as psychophysical performance.

Finally, we ran a feasibility experiment testing a subset of N = 5 participants with *vision-only* events, where the visual stimulus onset was delivered by keypress performed by the experimenter (average number of trials ± 1 standard deviation per participant: 13±0.5). The subject reported silently the task as in the main experiment. Also in this condition the timing of the visual presentation was jittered relative to the TR; circular statistics verified that the temporal distribution of events was homogeneous in the TR (Rayleigh test: p = 0.24).

### MRI scanning

Scanning was performed on a MR950 7T whole body MRI system (GE Healthcare, Milwaukee, WI, USA) equipped with a 2-channel transmit coil driven in quadrature mode, a 32-channel receive coil (Nova Medical, Wilmington, MA, USA) and a high-performance gradient system (50 mT/m maximum amplitude and 200 mT/m/ms slew rate).

For twelve participants, the anatomical images were acquired at 1 mm isotropic resolution using a T1-weighted magnetization-prepared Fast Spoiled Gradient Echo (FSPGR) sequence with the following parameters: TR = 6 ms, TE = 2.2 ms, flip angle = 12 deg, receiver Bandwidth = 50 kHz, TI = 450 ms, ASSET = 2. For the remaining 9 participants, high-resolution anatomical images with 0.8 mm isotropic resolution were acquired with a modified magnetization-prepared rapid gradient echo sequence (MP2RAGE) with the following parameters: TR = 6.6 ms, TE = 2.5 ms, flip angle = 5 deg, receiver Bandwidth = 62.5 kHz, TI = 1000 ms and 3200 ms, ARC factor = 2.

Functional images were acquired with a gradient-echo EPI sequence with slices N = 46 (with an ascending interleaved order), Field of View = 192 mm, matrix size = 128×128, resulting in a spatial resolution of 1.5×1.5 mm^2^ in-plane, and slice thickness 1.4 mm with slice spacing = 0.1mm, TR = 3 s, TE = 23 ms, flip angle = 60 deg, receiver Bandwidth = 250 kHz, ASSET acceleration factor = 2, phase encoding direction: Anterior-to-Posterior. No resampling was performed during the reconstruction. For each EPI sequence, we acquired an additional volume with the reversed phase encoding direction (Posterior-to-Anterior), used for distortion correction (see pre-processing).

### Pre-processing of imaging data

Anatomical images were processed by a standard procedure for segmentation implemented in Freesurfer (recon-all). In addition, hemispheres were aligned to a template of the cortical surface (fsaverage) as well as to the left/right symmetric version of the same template (fsaverage_sym).

We used Brain Voyager with default settings to perform slice time correction and motion correction. Geometrical distortions were compensated by using EPI images with reversed phase encoding direction (Brain Voyager COPE plug-in). The pre-processed images were aligned to each participant’s anatomical image using a boundary-based registration algorithm (Freesurfer “*bbregister”* algorithm) and projected to the cortical surface of each hemisphere. All analyses were conducted on data in the individual subject space, resampled to a resolution of 3×3×3 mm. For each run, we removed the first 6 TRs (to allow the MR signal to reach steady-state), then estimated the temporal linear trend and subtracted it from the fMRI time-course. Time-courses were averaged within each ROI (see below) and converted into percentage signal change by normalizing them to the mean BOLD.

## QUANTIFICATION AND STATISTICAL ANALYSIS

### Definition of Regions of Interest (ROI)

For each participant, we defined three ROIs (bilateral V1, left M1 and left S1) in the 3D native space of each participant. V1 was defined using the responses to the vertical and horizontal meridians from the retinotopic mapping scans and included the cortical representation of the stimulus area, in both hemispheres of about 7.3 cm^3^. The M1 and S1 ROIs were defined based on the pattern of BOLD responses to keypress associated with the perceptual decision that generated two separate foci located rostrally and caudally of the left Rolandic sulcus, consistent with M1 and Brodmann Area 3a. The M1 and S1 ROIs comprised on average 5.8 cm^3^ of cortex.

### Evaluation of fMRI Activity

We used the Finite Impulse Response deconvolution approach for estimating the Haemodynamic Response Function (HRF) over a window of 7 TRs following each event (an “event” being a visual stimulus presentation, a voluntary keypress or both). This analysis approach does not assume a particular shape of the haemodynamic response but, rather, estimates it directly by modelling each of the 7 TRs after the events as a separate GLM predictor and thereby assigning each with a beta-weight^70^. Given that visual stimuli and the keypress that started the trial were separated by less than 350 ms, we aligned the predictor with the TR that included the keypress (Figure S2A shows V1 BOLD responses computed after aligning to the visual stimulus instead: compare Figure S2A with Figure 2A in the main text). In each ROI, seven beta values per each event type were estimated with GLM (e.g. for the main analyses, we considered four types of events corresponding to the 4 visuo-motor delays, making 28 predictors to which we added a constant term); the average goodness of fit was 20±2% (average number of timepoints: 629).

Statistical comparisons were performed after extracting GLM beta-weights in individual participants and in each of our *a priori* defined ROIs.

We assessed pairwise correlations between M1 and V1 in the aggregate observer data, after regressing out the event-related modulations, i.e. using the residuals of the GLM model. Specifically, we performed 4 steps: 1) compute the predicted time-course given the estimated GLM weights for each run; 2) subtract it from the observed time-course to obtain the residuals; 3) concatenate residuals from all runs and all participants; 4) compute the Pearson’s correlation between residuals in the two ROIs. Correlation coefficients were then averaged in the −3 to 12 s interval around vision and action events, separately for the four visuomotor delays. Correlation differences across delays were assessed by bootstrap (10000 repetitions with replacement, one-tailed test), repeating step #4 10000 times for each delay. We complemented this aggregate subject analysis with a more standard repeated measures approach, where correlations were computed for each participant and visuomotor delay. Correlation differences across delays were then assessed with paired t-tests.

GLM beta weights at the TRs of interest (the peak of haemodynamic response at TR2 for Figure 1C and S1D, or the integral of beta weights (divided by time) from TR1 to TR4 for Figure 3 and S1D) were also visualized over a template of the cortical surface. These were estimated after projecting the pre-processed BOLD timecourses from each participant on a common template, concatenating them and then applying the GLM analysis. For Figure 1C and S1D-E, data from the two hemispheres were analysed separately; for Figure 3, a mirror-symmetric template of the cortical surface was used and the concatenated BOLD timecourses were averaged across hemispheres before the GLM analysis. Except for Figure 3A, activity maps were masked for statistical significance (p*FDR* < 0.05, Figure S1E reports uncorrected p-values). Eccentricities in primary visual cortex were defined according to the Benson template for fsaverage^71^.

## KEY RESOURCES TABLE

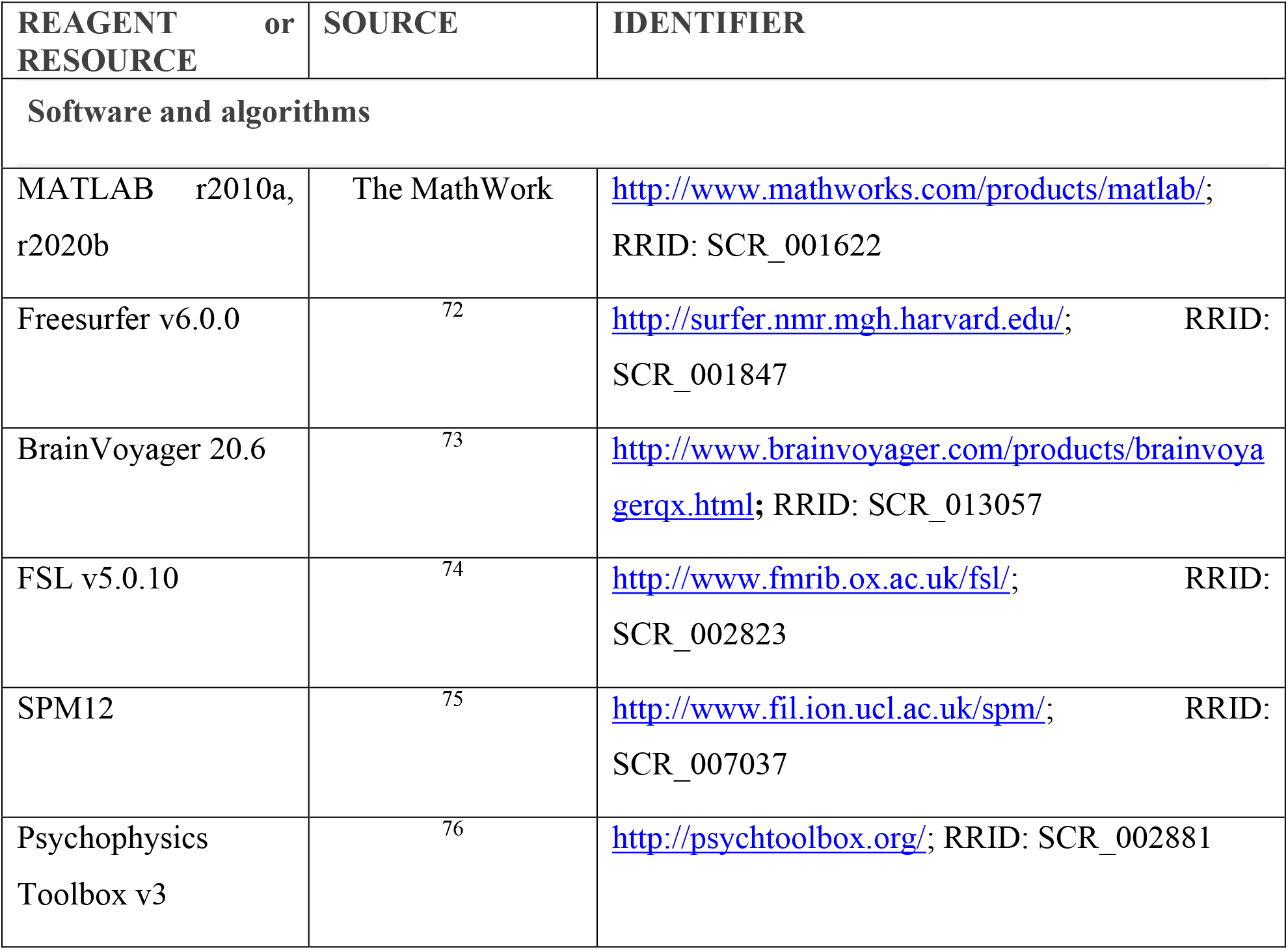

