## Supplementary Figure for "Predictive visuo-motor communication through neural oscillations"

### Supplemental figures

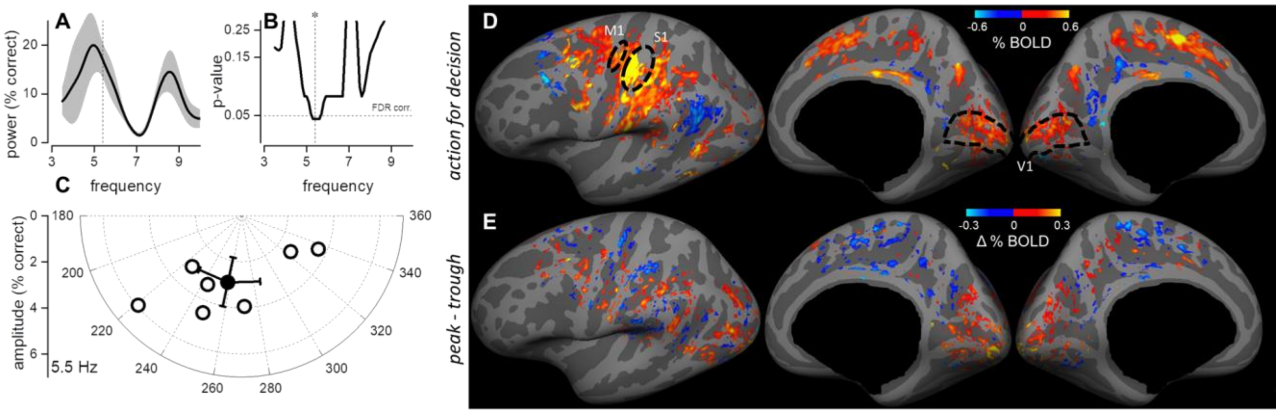


**Figure S1 related to Figure 1. Behavioural and BOLD responses**

**A-C**: consistency of perceptual oscillation across participants, obtained by fitting each participant's dataset with sinusoidal curves. **A:** Power and s.e.m averaged across participants as a function of frequencies between 3.5 and 10 Hz. **B:** FDR-corrected statistical significance, derived from Hotelling $T^2$ distribution across participants, which is maximal at 5.5 Hz, very similar to the best fit for the mean power across participants. **C.** Amplitude and phase of the best fitting of a sinusoidal function at 5.5 Hz in each participant, clearly clustered around 260°. The filled circle plots mean amplitude and phase with s.e.m.

**D**: Maps of the BOLD response to *action-only* events, measured in an independent set of trials where participants performed the 2AFC task. The activity is related to the keypress reporting the perceptual decision (separated by at least 15s from any *vision and action* event). BOLD activity is estimated as the peak of the haemodynamic, at 6s, and masked at 0.05 significance after FDR correction. Activity around the central sulcus of the left hemisphere defined the M1 and S1 ROI (in each participant's volume; black lines mark the approximate projection of these ROIs). Other foci of activity included the early visual cortex, which was strongly and unexpectedly modulated with *action-only* events, putative area V6 in the parieto-occipital[S1, S2] sulcus as well as portions of the cingulate cortex[S2], both similarly active in response to *vision and action* events (compare with Figure 1C of the main text).

**E**: Contrast of the BOLD response to *vision and action* events at SOAs corresponding to the peak vs. trough delays of psychophysical performance, masked at 0.05 significance uncorrected and computed as the integral from 3 to 12 s of the haemodynamic response, divided by time. Positive modulations clustered in the occipital lobe; there was no clear positive or negative modulation in the other foci of activity for the vision and action events (see Figure 1C in the main text): neither putative area V6 in the parieto-occipital sulcus, nor the cingulate sulcus visual area or the somatosensory and motor areas show consistent clustering of contrast values.

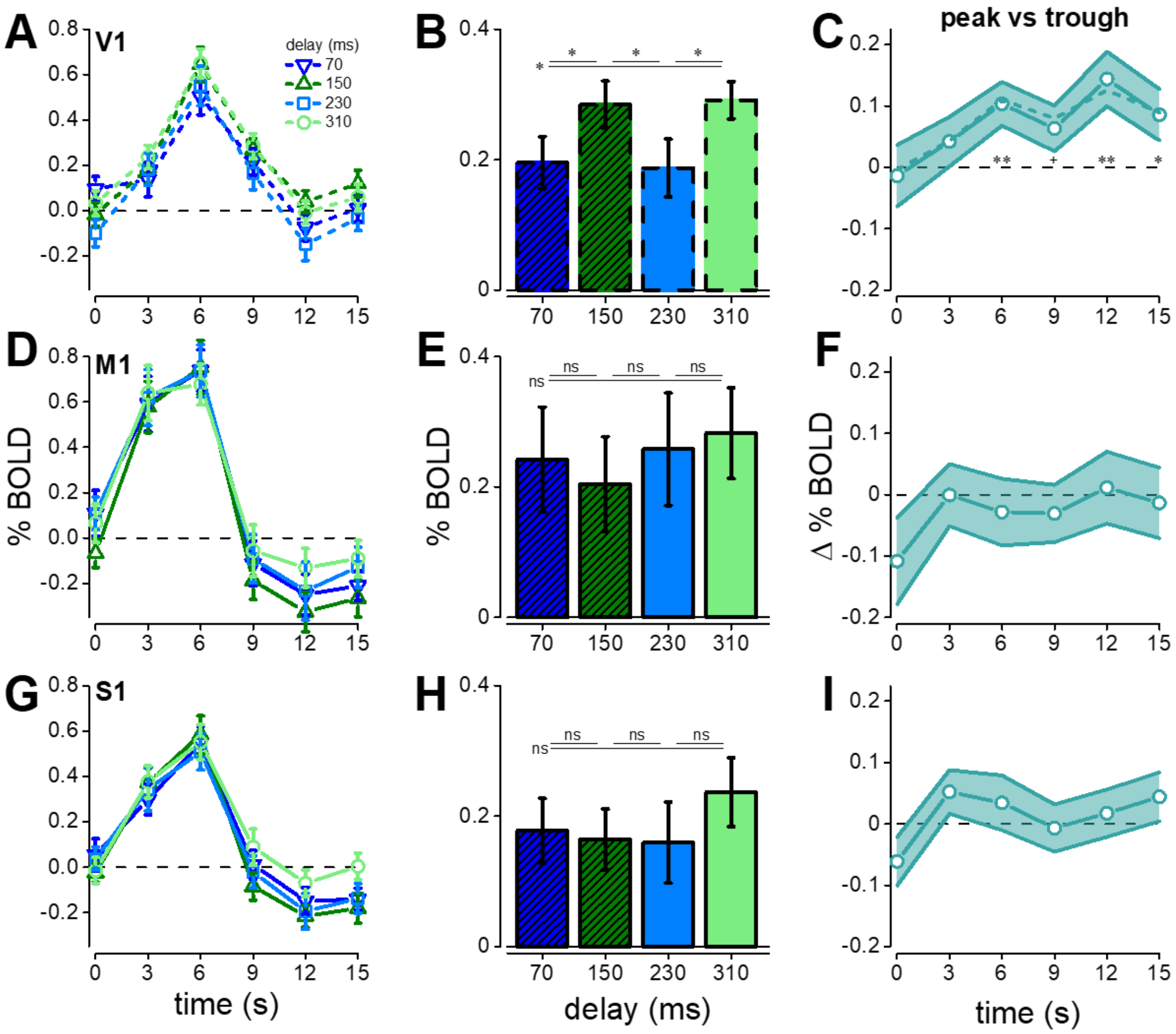


**Figure S2 related to Figure 2. V1, M1 and S1 BOLD responses to *vision and action* events.**

**A-B**. same as B-C in Figure 2 of the main text, but with GLM beta weights for the V1 BOLD response estimated after aligning data to the onset of the visual stimulus, rather than to the keypress action. Asterisks in B mark statistical significance (0.05 > * > 0.01) of post-hoc paired one-tailed t-tests (Bonferroni-Holm corrected for multiple comparisons) comparing pairs of visuo-motor delays: 70 ms vs 150 ms: $t(16) = 2.69$, $p = 0.032$; 150 ms vs 230 ms: $t(16) = 2.64$, $p = 0.032$; 230 ms vs 310 ms: $t(16) = 2.47$, $p = 0.026$; 70 ms vs 310 ms: $t(16) = 2.45$, $p = 0.025$).

**C**. Modulation of the V1 response to vision and action events associated with the peak vs. trough delays of psychophysical performance, computed for each TR following the keypress action (continuous lines) or the stimulus (dashed lines). For the continuous line, statistical significance is for TRs: 0 s: $t(16) = 0.27$, $p = 0.60$; 3 s: $t(16) = 1.04$, $p = 0.15$; 6 s: $t(16) = 2.90$, $p = 0.005$; 9 s: $t(16) = 1.70$, $p = 0.054$; 12 s: $t(16) = 3.23$, $p = 0.002$; 15 s: $t(16) = 2.04$, $p = 0.028$ (p: 0.06 > + > 0.05 > * > 0.01 > ** > 0.001).

**D-I**. BOLD response for vision and action events associated with the peak vs. trough delays of psychophysical performance in M1 (D, E, F) or S1 (G, H, I). No comparisons reach significant threshold ($p<0.16$, uncorrected).

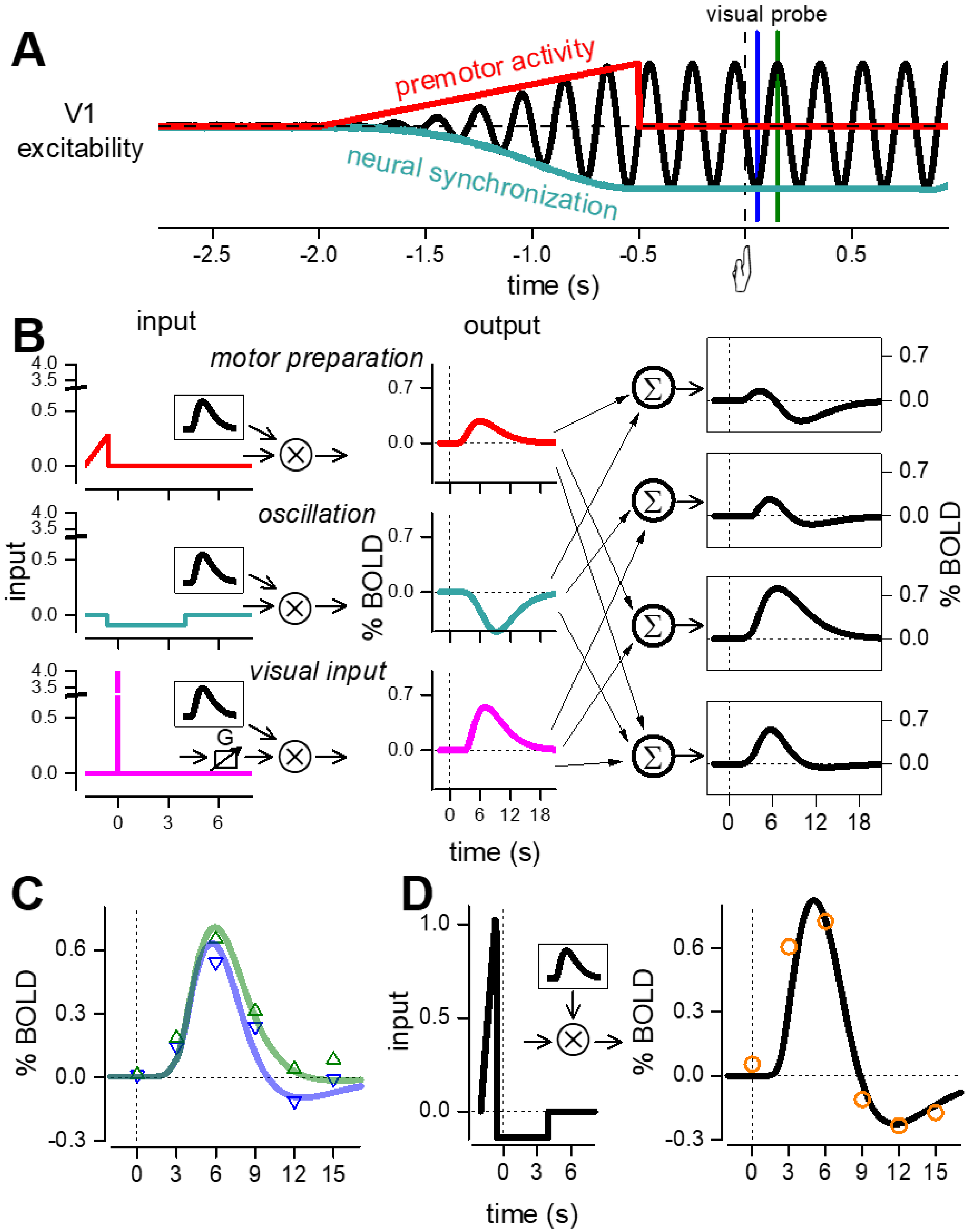


**Figure S3 related to Figure 4. Simulation of the V1 and M1 BOLD response and their functional connectivity.**

**A**. We assume that the neural activity underlying BOLD responses to *vision and action* events can be described with the following components: 1) a ramp preceding the action and representing premotor activity, e.g. in the form of a readiness potential (red curve; onset −2 s, offset −0.55 s from action onset); and 2) an oscillatory signal, generated by the phase synchronization of endogenous rhythms around 5.4 Hz (black curve), which reaches maximum amplitude before the action onset and persists for several seconds after (from −0.55 s to +4 s; schematically represented by the black curve). We further assume that V1 response gain is dynamically modulated by the oscillatory signal, implying that responses to identical visual stimuli are more enhanced when presented at the peak of the oscillation (green line) then when presented at the trough (blue line).

**B,C,D**. To simulate the BOLD response, the above signals are convolved with the standard model of the haemodynamic response given by:

$$IRF(t) = \frac{\left(\frac{t-\delta}{\tau}\right)^{(n-1)} e^{-\left(\frac{t-\delta}{\tau}\right)}}{\tau(n-1)!} \qquad \text{[eq. S1]}$$

With decay time $\tau = 2s$, delay $\delta = 3s$, and order $n = 3$.

To keep model complexity low, the visual response is modelled by a delta function, and pre-motor activity is the same as in A. The synchronization of endogenous oscillations has been simulated with a negative constant signal (blue-green curve), given previous evidence that oscillatory phase coherence of endogenous rhythms is associated with negative BOLD[S3, S4]. The visual response amplitude changes with delay to simulate the action of gain (G) set by the synchronization.

Different combinations of the three inputs, after convolution with the *IRF(t)*, fit the different BOLD responses. The amplitude of the ramp was set to 0.27 by best fit ($R^2$ =0.74) of the V1 BOLD response to *action-only* events (see maps in Figure S1 and Figure 4 red curve).

The amplitude of the delta function was set to 4 by best fitting ($R^2$=0.91) the response of the *vision-only* response (Figure 2A, pink curve). These values were then used as fixed parameters in the fit of the BOLD response in V1 as a function of delay.

The timing of the offset of the ramp and onset of the negative signals were set to −550 ms by best fitting of the data of the M1 responses as shown in D. The best fit of M1 ($R^2$=0.89) was obtained with ramp amplitude equal to 1.10 and negative component equal to 0.13.

Any linear combination of the BOLD response to the ramp (first row in panel B) and delta functions (third row in panel B) will be inadequate to fit the M1 or V1 BOLD response to *vision and action* events, given the slow decay of the sum (third panel of the left column, in B) and the lack of a negative lobe. However, the linear combination of these two signals is necessary to reproduce the fast growth of the visual response in the first two TR (6 s) to *vision and action* compared to the *vision-only* events (compare dashed black curve with blue curve in Figure 4C). To model the response to *vision and action* of V1, it was necessary to add the negative signal. The result adequately fit ($R^2$=0.92) the response to *vision and action* for the trough delays (blue curve or Figure 4C, blue curve) using an amplitude of the constant signal equal to −0.08. To fit adequately the peak delay responses, which differ from the trough especially for the reduced negative lobe, it is necessary to change the ratio between amplitude of response and amplitude of negative BOLD.

A good fit was obtained by modulating the gain of the visual response by a factor of 1.3 (Figure 4C). The alternative strategy of reducing the negative signal amplitude also allows fitting of the response to trough delays, obtaining an $R^2$ value of 0.88, lower than increasing the gain equal to 0.92 (Figure S3,C). In addition, a reduction of negative signal is difficult to explain given that this signal is present before the presentation of the visual stimulus. Both gain modulation and constant signal have been shown to be important to simulate how LFP in sensory cortex changes with synchronization of rhythms[S5].

Gain modulation may also be crucial to explain the variation of functional connectivity between M1 and V1 with delay. Single cell recordings have demonstrated that correlation between neurons can be modelled considering that the total variance of the discharge is weighted by the gain variance[S6]. To test if similar mechanisms can be applied for BOLD functional connectivity, we simulated the modulation of V1 visual response by M1 noise given by:

$$V1_{out}(t) = \alpha \cdot \left(IRF(t) \otimes S(t)\right) \cdot N_{M1}(t) + N_{V1}(t) \qquad \text{[eq. S2]}$$

and

$$M1(t) = N_{M1}(t) \qquad \text{[eq. S3]}$$

where $IRF(t)$ is the neuronal impulse response function estimated by the Gamma function of equation S1 with a time decay $\tau$ =80 ms, $\delta = 15ms$ and order $n$ of 2; $N_{M1}$and $N_{V1}$are white noise of specific amplitude and free parameter variance. $S$ is a series of about 16 events randomly positioned in a time interval of 300 s.

We applied to $V1_{out}$ the same GLM adopting the same finite IRF approach used for our analyses with 60 equi-spaced regressors in time separated by 10 ms. We verified that the GLM produced unbiased estimates of the visual response in V1 (despite the presence of a non-linearity), and that the residuals did not contain IRF(t) information. The correlation between V1 and M1 (for time window containing the first 200 ms of IRF duration) increased as a function of gain increase ($\alpha$). Enhancing excitability in V1 by a factor of 1.3 (the same factor used to model the visual responses in Figure 4C) increased the V1-M1 correlation by about 1.27. The parameter to achieve a modulation that fits well the data of Figure 2E were $N_{V1}$, a white noise in the range [0:0.4], $N_{M1}$is a motor noise in the range [0:0.1], $\alpha$ is equal to 2. Convolving $V1_{out}$ and *M1* of equations S2 and S3 with the haemodynamic function of equation S1, and repeating the procedure increased correlation but did not change the modulation with visuo-motor delay. The simulation reinforces the concept that gain modulation may be a key mechanism for functional coupling of distal brain regions.
